# Environmental rhythms orchestrate neural activity at multiple stages of processing during memory encoding: Evidence from event-related potentials

**DOI:** 10.1101/2020.06.02.129379

**Authors:** Paige Hickey, Annie Barnett-Young, Aniruddh D. Patel, Elizabeth Race

## Abstract

Accumulating evidence suggests that rhythmic temporal structures in the environment influence memory formation. For example, stimuli that appear in synchrony with the beat of background, environmental rhythms are better remembered than stimuli that appear out-of-synchrony with the beat. This rhythmic modulation of memory has been linked to entrained neural oscillations which are proposed to act as a mechanism of selective attention by amplifying early sensory responses to events that coincide with the beat. The current study aimed to further test this hypothesis by using event-related potentials (ERPs) to investigate the locus of stimulus processing at which rhythm temporal cues operate in the service of memory formation. Participants incidentally encoded a series of visual objects while passively listening to background, instrumental music with a steady beat. Objects either appeared in-synchrony or out-of-synchrony with the background beat. Participants were then given a surprise subsequent memory test (in silence). The timing of stimulus presentation during encoding (in-synchrony or out-of-synchrony with the background beat) influenced canonical ERPs associated with post-perceptual selection and orienting attention in time. Importantly, post-perceptual ERPs also differed according to whether or not participants demonstrated a mnemonic benefit for in-synchrony compared to out-of-synchrony stimuli, and were related to the magnitude of the rhythmic modulation of memory across participants. These results support two prominent theories in the field, the Dynamic Attending Theory and the Oscillation Selection Hypothesis, which propose that neural responses to rhythm act as a core mechanism of selective attention that optimize processing at specific moments in time. Furthermore, they reveal that in addition to acting as a mechanism of early attentional selection, rhythm influences later, post-perceptual cognitive processes as events are transformed into memory.

## Introduction

Rhythmic temporal cues abound in our environment. A large body of research has demonstrated that exposure to environmental rhythms influences perception and action by enhancing processing at specific moments of time that align with the rhythmic beat [1-3]. More recently, there has been growing interest in how environmental rhythms influence higher-order cognitive processes such as memory formation [4-7]. In a series of studies, Johndro and colleagues found that the timing of individual events within a rhythmic temporal stream influenced memory formation [8]. Specifically, stimuli presented in alignment with the timing of a background rhythm (on-beat) were better remembered in subsequent tests of memory than stimuli presented out-of-alignment (off-beat). This rhythmic modulation of memory (RMM) occurred even when rhythmic temporal cues were task-irrelevant and presented in a different modality (auditory) than the target stimuli (visual), suggesting that rhythmic cues guide domain-general attentional resources to specific moments in time and enhance the processing of information that occurs in alignment with the beat [9-11]. Importantly, these results provide evidence that orienting attention to specific moments in time, like orienting attention to specific spatial locations or object features, plays a critical role in selecting what environmental information will be effectively encoded into long-term memory.

Electrophysiological evidence further supports the notion that dynamic fluctuations in attention underlie rhythmic modulations of memory encoding. It is well established that neural activity synchronizes to the periodicity of external rhythms, evident in increased power and phase alignment of neural oscillations at the same frequency as the external rhythmic stream [1,12,13]. The entrainment of low-frequency neural oscillations, particularly in the delta range, has been shown to dynamically modulate fluctuations in cortical excitability, such that high excitability states align with the temporal pattern of the rhythmic stream [11,14]. This rhythmic shift in neural excitability has been proposed to act as a core mechanism of selective attention that optimizes stimulus processing at specific moments in time (in phase with the rhythmic beat) [14-16]. Support for this Oscillation Selection Hypothesis comes from numerous studies demonstrating enhanced perceptual processing of stimuli which appear at rhythmically-predicted moments in time [1,14,15,17-21]. Recent evidence suggests that neural entrainment to low-frequency rhythm also modulates higher-order cognitive processing and the encoding of events into long-term memory [22]. In this study, participants incidentally encoded a series of visual objects while passively listening to background, instrumental music with a steady beat. Like the prior behavioral study by Johndro and colleagues, objects either appeared in-synchrony or out-of-synchrony with the background beat [8]. Participants were then given a surprise subsequent memory test (in silence). Results revealed significant neural tracking of the musical beat at encoding, evident in increased electrophysiological power and inter-trial phase coherence at the perceived beat frequency (1.25 Hz). Importantly, enhanced neural tracking of the background rhythm at encoding was associated with superior subsequent memory for in-synchrony compared to out-of-synchrony objects during a later memory test, revealing that neural responses to rhythm influence memory formation at specific moments in time. An important outstanding question is the specific mechanism by which external rhythms influence stimulus processing en route to memory formation.

One possibility is that rhythmic temporal cues influence memory encoding by modulating early sensory or perceptual processing. According to the Oscillation Selection Hypothesis, neural entrainment to external rhythms acts as an early sensory gain control mechanism that amplifies stimulus-driven neural activity at specific moments in time (in phase with the rhythm) [2,14,15]. Indeed, selective attention is known to modulate the amplitude of early stimulus-evoked potentials associated with initial sensory/perceptual processing [23-25]. For example, a well-established finding in the spatial attention literature is that directing attention to the location where a stimulus will appear enhances the amplitude of visual ERP components, such as the N1 associated with initial stimulus processing [23,26]. These early amplitude modulations have been interpreted as reflecting gain control or amplification mechanisms within early visual pathways that enhance stimulus perception [23-24]. In a recent study, Escoffier and colleagues extended these results to the temporal domain and found that the presentation of visual stimuli at temporally-predicted moments (in synchrony with a background auditory beat) enhances the amplitude of the N1 component over posterior electrode sites [27]. By this view, rhythm could influence memory encoding by enhancing early perceptual processing of stimuli that appear in synchrony with the beat.

In addition to influencing early sensory-perceptual processing, rhythmic temporal cues could also influence memory encoding by modulating later, post-perceptual stages of stimulus processing. It is well known that attention can bias stimulus processing at multiple stages of information processing [23-25,29-30]. Prior studies have shown later ERP components associated with post-perceptual stimulus identification and evaluation (e.g., N2, P3) [31-35] are sensitive to temporal orienting cues that direct attention to particular moments in time [30,36,37]. It has also been suggested that manipulations of temporal attention may have their greatest effect on these later, post-perceptual stages of information processing [30,34,38]. In support of this proposal, effects of temporal orienting on early visual ERP components are not always observed and have been shown to depend on the perceptual demands of the task and the nature of the temporal orienting cues [19,30,38-42]. Thus, the effect of rhythm on memory encoding could primarily reflect changes in later, post-perceptual stages of information processing.

Findings from the memory literature provide additional support for the possibility that later stages of processing may be the locus of rhythmic effects on stimulus encoding. Electrophysiological studies have identified a slower, positive-going waveform that occurs ∼500ms after stimulus presentation and predicts whether or not a stimulus will be later remembered or forgotten [43-45]. This later positive complex (LPC), which is often greatest over more anterior electrode sites, is enhanced in task contexts that strengthen memory encoding (e.g., deep vs. shallow processing, semantic vs. non-semantic classification) and has been interpreted as reflecting more elaborative, higher-level cognitive processing that leads to the formation of a more durable memory trace [45,46-48]. Although there has been little research into the effects of temporal orienting on electrophysiological responses occurring during this later time window, modulations of this more sustained frontal waveform may be particularly relevant to effective memory encoding.

To date, only one prior study has attempted to identify the locus of rhythmic effects on memory encoding using ERPs [7]. In this study, participants explicitly encoded visual targets that occurred in either a rhythmic temporal structure (separated by a regular interstimulus interval) or an arrhythmic temporal structure (separated by an irregular interstimulus interval). Rhythmic presentation had a significant effect on memory encoding, with superior subsequent memory for stimuli presented in the rhythmic compared to arrhythmic context. Rhythmic presentation also influenced stimulus-evoked neural responses at encoding, modulating both the early visual N1 component occurring ∼200ms after stimulus presentation, as well as a late positive complex (LPC) starting ∼400ms after stimulus presentation (post-perceptual N2/P3 amplitudes were not investigated). These results suggest that rhythmic temporal cues can influence both early perceptual as well as later cognitive stages of stimulus processing. However, modulation of both early and later ERP components was negatively associated with the effect of rhythm on memory performance, raising questions about the behavioral relevance of these evoked neural responses to rhythm. In addition, it is important to note that this study was unable to test whether evoked responses differ according to the timing of stimulus presentation within a rhythmic temporal stream (e.g., on-beat vs. off-beat). Such a comparison is critical in order to test the hypothesis that rhythmic temporal cues influence memory encoding by guiding attention to expected moments in time.

The current study leveraged the high temporal resolution of ERPs to explore how the timing of stimulus presentation within a rhythmic temporal stream influences different stages of stimulus processing during memory encoding. During EEG recording, participants incidentally encoded visual stimuli that appeared either in synchrony (on-beat) or out-of-synchrony (off-beat) with a background, auditory rhythm. Previously, we demonstrated that while participants generally demonstrate superior subsequent memory for visual stimuli that appear on-beat compared to off-beat at encoding, there is substantial individual variability in the effect of rhythm on memory performance [8,22]. Importantly, while some individuals demonstrate superior subsequent memory for on-beat compared to off-beat stimuli, others do not, and this variability in the rhythmic modulation of memory is closely linked to individual differences in the fidelity by which external rhythms are represented in the brain (neural entrainment). The current study built upon these findings to examine whether the effect of rhythm on memory performance is also related to stimulus-evoked neural responses to rhythm. We first examined whether the timing of visual stimulus presentation within the auditory rhythm (on-beat vs. off-beat) influenced attention-related ERPs during early (perceptual) stages of processing and/or later (post-perceptual) stages of processing. If the rhythmic modulation of memory reflects attentional selection early in the processing stream (e.g., perceptual selection), the amplitudes of early perceptual ERPs (e.g., N1) should differ for on-beat and off-beat stimuli. If the rhythmic modulation of memory reflects attentional selection after perceptual processing is complete (e.g., post-perceptual selection) or changes in higher-order stimulus processing (e.g., cognitive control), later ERPs (N2, P3, LPC) should differ for on-beat and off-beat stimuli. Next, we aimed to more directly examine the relationship between stimulus-evoked neural responses and the effects of rhythm on memory formation by exploring whether ERPs differ according to whether or not individuals demonstrate a rhythmic modulation of memory effect (e.g., better subsequent memory for on-beat than off-beat stimuli). We also tested whether individual differences in stimulus-evoked neural activity predict the degree to which rhythm modulates memory encoding across participants.

## Materials and Methods

### Participants

A sample of 36 participants (12 male, 24 female) between 18-31 years of age (*M =* 23, *SD =* 3.32) were recruited from Tufts University and the surrounding community. Participants were either compensated $15/hour or received course credit for participation. All participants were right-handed, fluent English speakers, and had normal or corrected-to-normal eyesight and hearing. Additionally, participants were screened for history of neurological illness, brain injury, substance use, and psychiatric diagnosis. All participants provided written informed consent in accordance with the procedures from the Institutional Review Board at Tufts University.

### Procedure

This study is a new analysis of the EEG data collected as part of the experiment previously reported by Hickey and colleagues [22]. Participants performed an incidental encoding task in which they were presented 120 images (60 animate, 60 inanimate) from the Multilingual Picture Database (MultiPic) [49] in the presence of background music (Fig 1). The background music was created in Garage Band and was designed to contain a steady beat with a tempo of 75bpm or 1.25 Hz (inter-beat interval = 800ms) and a 4/4 metrical structure. The music did not contain words or lyrics and was designed to sound naturalistic, without being highly complex in melody, rhythm, or texture. Each image was presented in the center of the screen for 750ms and participants were instructed to make a semantic classification about whether the image was living (animate) or non-living (inanimate) as quickly and accurately as possible. The timing between images was jittered (*M* = 6.4s, *SD* = 1.25s) with interstimulus intervals ranging from 3.75s to 7.75s. Critically, images were presented either in-synchrony with the background rhythm (on-beat) or 250ms prior to the beat (off-beat). The presentation order of the images was semi-randomized to ensure that equal numbers of animate/inanimate and on/off beat images were presented in each third of the experiment and that no more than six images of the same condition (on/off) appeared consecutively. EEG was recorded continuously throughout the incidental encoding period, which lasted approximately 13 minutes.

**Fig 1.**
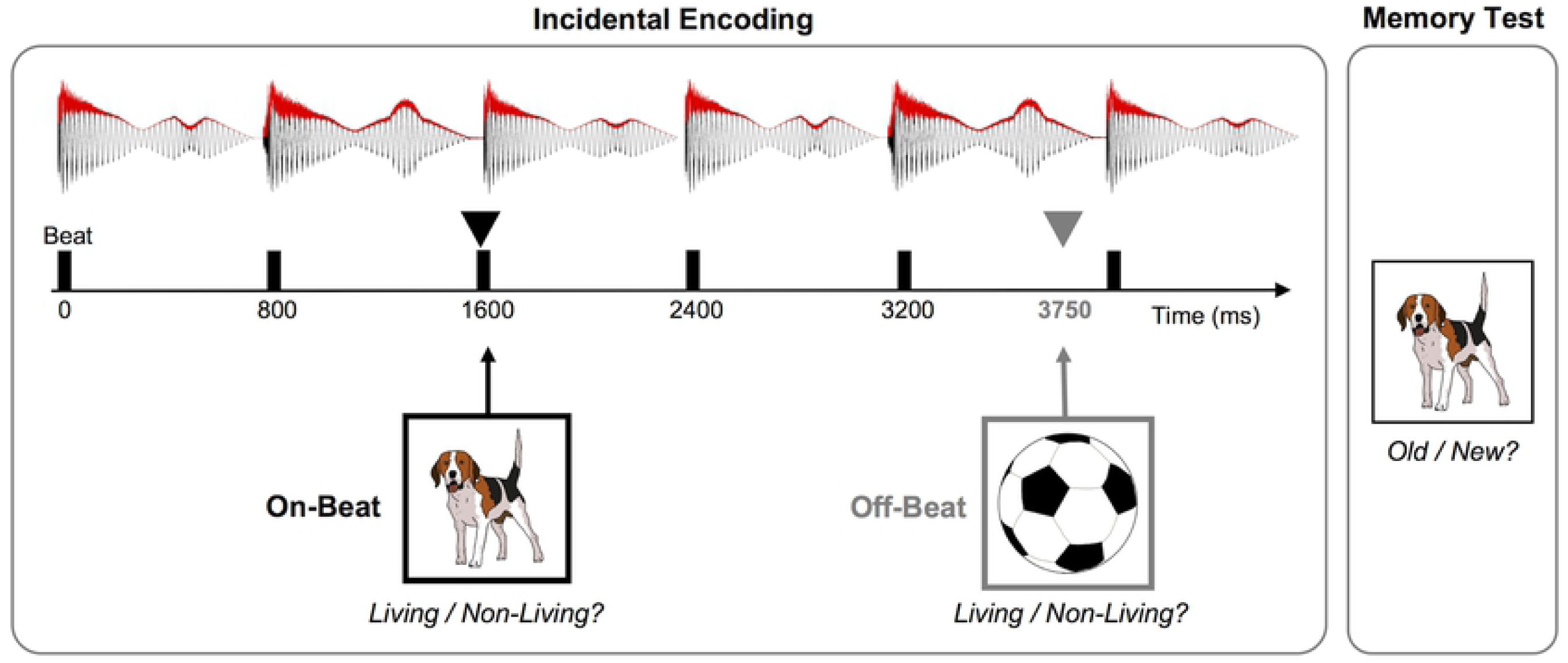
Behavioral paradigm. During incidental encoding, participants were shown a series of full color or black and white pictures of objects in the context of background, rhythmic music and made a semantic decision (living / non-living) about each object. Pictures remained on the screen for 750ms and appeared either in-synchrony (on-beat) or out-of-synchrony (off-beat) with the background rhythm (off-beat pictures were presented 250ms prior to the beat). Participants were then given a surprise memory test in silence in which they were shown pictures of objects that had been previously viewed during the encoding block and novel objects and made a recognition memory decision for each (old / new). The top of the figure displays an example of the musical waveform (black) and amplitude envelope (red) with vertical black lines indicating the timing of the underlying beat (800ms ISI, or 1.25 Hz) The lower portion of the figure shows example images with a thin arrow above each image showing the time of its onset.

After the incidental encoding period, participants were immediately given a self-paced surprise subsequent memory test (in silence). During the memory test, all 120 images from the encoding period and 60 new lure images were presented one at a time and participants decided whether they had previously seen the image in the first part of the experiment by making a yes/no button press on a keyboard. After making their memory decision, participants rated how confident they were in their decision (low or high). EEG was not recorded during the memory test.

### EEG Recording and Preprocessing

EEG was recorded using a BioSemi Active-Two amplifier system (Biosemi, Amsterdam, Netherlands) from 32 Ag/AgCl scalp electrodes and two reference electrodes place on the left and right mastoids. Two additional electrodes were also placed around the eyes to monitor eye movements. EEG pre-processing was completed using EEGlab and custom MATLAB scripts. All EEG signals recorded with a sampling rate of 1024 Hz and subsequently down-sampled to 512 Hz. EEG recordings from the 32 scalp electrodes were referenced to the average of the two mastoid electrodes and filtered using a 0.1 Hz high pass and 50 Hz low pass filter. Independent components analysis was then used to remove artifacts in the signal consistent with eye-blinks and muscle movements. Finally, the continuous signal was epoched from −2000ms to 2000ms surrounding visual stimulus presentation. Each epoch only contained one visual stimulus and epochs containing artifacts were manually rejected. Epochs were then baselined from −100ms to 0ms. Event related potentials were separately generated for on-beat and off-beat stimuli by averaging trials for each condition within each participant. Average event related potentials were also generated for the comparison of individuals who demonstrated greater memory for on-beat compared to off-beat stimuli (rhythmic modulation of memory; RMM group) and individuals who did not (no-RMM group).

### EEG Data Analysis

EEG analysis focused on the early N1 component associated with perceptual processing, as well as later ERP components associated with post-perceptual processing (N2, P3) and subsequent memory (LPC). For the N1, analysis time windows and posterior electrodes of interest were selected based on prior studies investigating rhythmic effects on perception [7,27,34]. Specifically, the peak of the N1 was determined based on the average waveform over bilateral posterior electrodes (O1, O2) and mean amplitudes were calculated and analyzed across a 20ms latency window around the peak (197-217ms). Slightly larger analysis time windows were selected for the later components following the approach used in prior studies investigating temporal cuing effects on stimulus processing [30,37] and subsequent memory [7,44]. Mean amplitudes for the N2 were calculated across a 40ms window around its peak (281-321ms) [30] and mean amplitudes for the P3 were calculated across a 100ms window around its peak (350-450ms) [37]. Mean amplitudes of the late positive complex (LPC) were calculated across a time window ranging from 450-750ms [7,44]. For the later components, analysis focused on more frontocentral electrodes (Cz, Fz) given the more fronto-central distribution of post-perceptual effects related to attention and mnemonic processing [30,34,44,47].

### Behavioral Data Analysis

Memory performance was measured by calculating the proportion of correctly identified old images (hits) and the proportion of incorrectly identified new images (false alarms) separately for on-beat and off-beat stimuli to calculate a measure of d-prime (d’) for each condition (on-beat, off-beat). Analysis of memory performance was collapsed over confidence ratings to increase power and to replicate prior work [8,22]. Trials that were inaccurately responded to during encoding or trials with a reaction time >2SD from the mean during encoding were excluded from the analysis. Rhythmic modulation of memory (RMM) index scores were generated by subtracting the d’ measure for off-beat trials from the d’ for on-beat trials. A summary of the behavioral results is provided in Table 1.

**Table 1.**
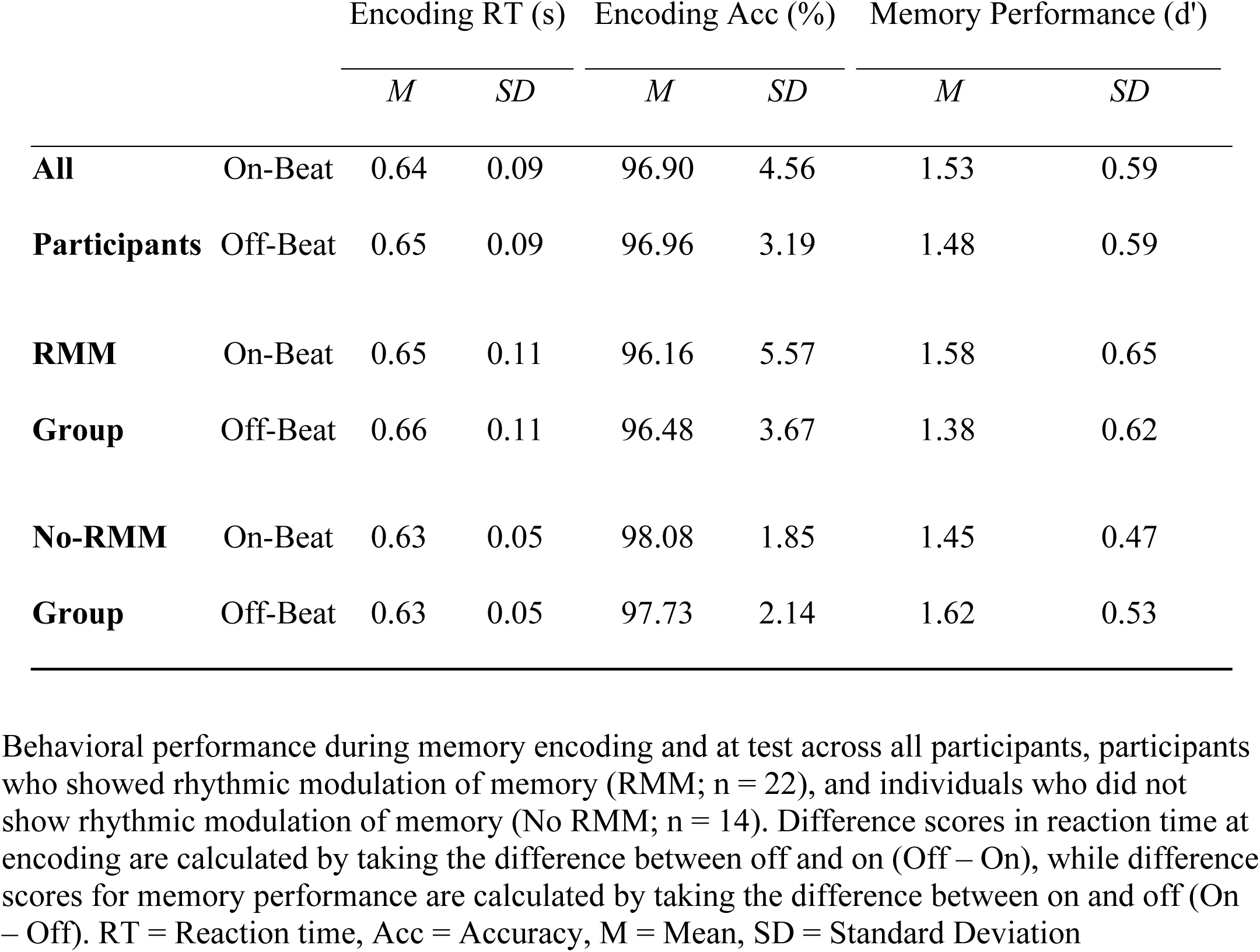
Behavioral Performance.

### Statistical Analysis

Separate statistical analyses were performed for each component/time window of interest. We first examined the effect of stimulus timing within the rhythmic temporal stream by comparing mean amplitudes for on-beat and off-beat trials. For the early N1, mean amplitudes were entered into a repeated-measures ANOVA with within-subject factors of timing (on-beat, off-beat) and electrode site (O1, O2). For the later components (N2, P3, LPC), mean amplitudes were entered into separate repeated-measures ANOVAs with within-subject factors of timing (on-beat, off-beat) and electrode site (Cz, Fz). Next, we investigated the effect of rhythmic timing on memory performance by investigating amplitude modulations in participants who demonstrated a rhythmic modulation of memory effect (RMM group; n = 22) and those who did not demonstrate a rhythmic modulation of memory effect (No-RMM group; n = 14). This analysis was motivated by the prior observation of significant individual differences in the effect of rhythm on neural activity and memory performance [22]. Pearson correlations and sequential multiple regressions were performed to further explore the relationship between stimulus-evoked ERP responses and the effect of rhythm on behavioral performance (rhythmic modulation of memory index) across individuals. All analyses were completed using SPSS version 25 and the PROCESS version 3.5 macro.

### Data Availability

The data that support the findings of this study are available online through Open Science Framework (DOI 10.17605/OSF.IO/WZC2G).

## Results

### Behavioral Results

Behavioral performance has been previously described by Hickey and colleagues (2020) [22] and is reported in Table 1. During encoding, reaction times were significantly faster for stimuli presented in-synchrony compared to out-of-synchrony with the background beat (*Z* = − 2.25, *p* = .01, one-tailed, *r* = .37), consistent with the proposal that rhythmic temporal cues orient attention to particular moments in time. During the subsequent memory test, memory performance was numerically greater for on-beat stimuli (*M* = 1.53, *SD* = .59) compared to off-beat stimuli (*M* = 1.48, *SD* = 0.59) across participants. However, substantial individual variability was present in the effect of rhythm on subsequent memory (RMM index range = −.54 to .49, *SD* = .23, c.f. Figure 3C of [22]), and the difference between on-beat and off-beat memory did not reach significance across participants (*t*(35) = 1.28, *p* = .21, one-tailed, *d* = .21). Previously, Hickey and colleagues (2020) found that variability in the effect of rhythm on memory performance was related to variability in participants’ neural responses to rhythm (neural entrainment), with better memory for stimuli presented on-beat versus off-beat at encoding (rhythmic modulation of memory) only observed in participants who demonstrated strong neural tracking of the beat [22]. This motivated separate analysis of individuals demonstrating a rhythmic modulation of memory effect (RMM; n = 22) and individuals who did not demonstrate a rhythmic modulation of memory effect (no-RMM; n = 14) in the present study. Behavioral data split by group is also presented in Table 1. In the RMM group, on-beat images were responded to faster than off-beat images at encoding (*Z* = −2.29, *p* = 0.02, *r* = 0.49) and were also better remembered than off-beat images in the subsequent memory test (*t*(21) = − 2.63, *p* = 0.016, *d* = 0.56). In contrast, the no-RMM group did not demonstrate a significant difference in RTs for on-beat versus off-beat images at encoding (*Z* = −0.79, *p* = 0.43, *r* = 0.21) and subsequent memory was reduced in the no-RMM group for on-beat compared to off-beat trials (*t*(13) = −4.02, *p* = 0.001, *d* = 0.15).

**Fig 2.**
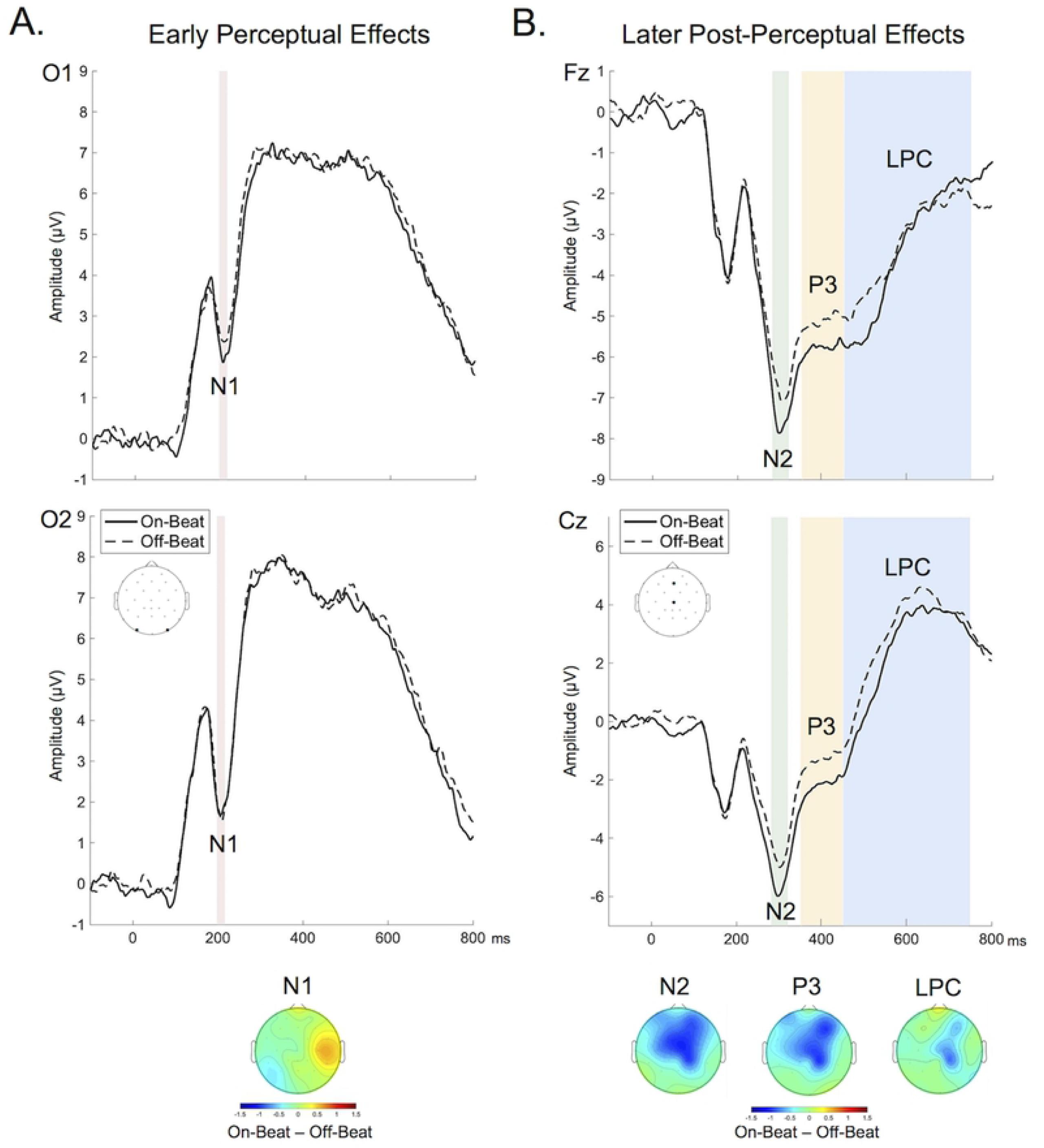
ERP waveforms for on-beat and off-beat trials. (A) ERP waveforms in posterior electrodes (O1, O2) were evaluated for early perceptual components. There was no significant difference in the early perceptual component (N1) between on-beat and off-beat trials. (B) ERP waveforms for on-beat and off-beat trials were plotted at electrodes Fz and Cz for post-perceptual components (N2, P3, LPC). The N2 and P3 components showed significant differences between on-beat and off-beat trials, but there was no difference in the LPC. Topography suggests that the greatest differences between on-beat and off-beat trials in the N2 and P3 component are present in fronto-central regions.

**Fig 3.**
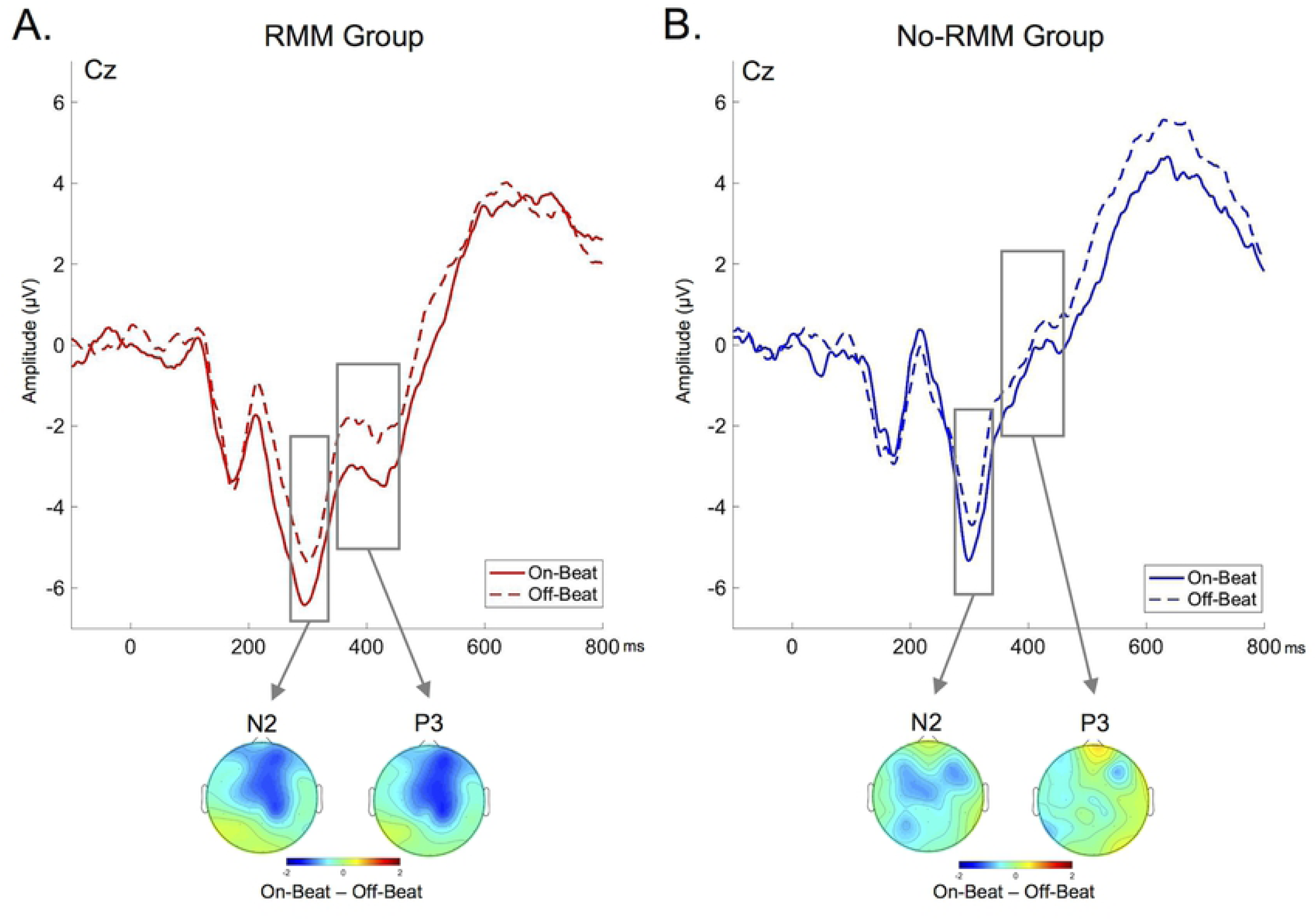
ERP waveforms for on-beat and off-beat trials time-locked to the onset of the visual stimulus at representative electrode Cz in each memory group. (A) RMM group, (B) No-RMM group. Scalp topography of the average signal differences (on-beat – off-beat) trials in each memory group during the N2 and P3 time window. Amplitude differences between on-beat and off-beat trials are present in the RMM group, particularly in frontal and central regions, but not in the No RMM group.

### EEG Results

#### Effects of stimulus timing

We first investigated whether the timing of visual stimulus presentation within the auditory rhythmic stream (on-beat vs. off-beat) influenced the amplitude of early perceptual (N1) or later post-perceptual (N2, P3, LPC) ERPs (Fig 2). Stimulus timing did not affect the amplitude of the early N1 (Fig 2A), with no main effect of stimulus timing, electrode, nor timing x electrode interaction (*p*s > .14). In contrast, the timing of stimulus presentation did affect the amplitude of the later N2 and the P3 components (Fig 2B). For the N2, there was a main effect of stimulus timing (*F*(1,35) = 9.39, *p* = .004, η^2^_p_ = .21), reflecting more negative amplitudes for on-beat compared to off-beat stimuli. There was also a main effect of electrode (*F*(1,35) = 12.84, *p* = .001, η^2^_p_ = .27), but no timing x electrode interaction (*F* < 1). For the P3 component, the main effect of stimulus timing approached significance (*F*(1,35) = 3.80, *p* = .06, η^2^_p_ = .10), with more positive amplitudes for off-beat compared to on-beat stimuli. There was also a main effect of electrode (*F*(1,35) = 34.69, *p* < .001, η^2^_p_ = .50), but no timing x electrode interaction (*F* < 1). Given that the effect of temporal orienting on P3 amplitudes is often largest over central electrode sites (Cz) [30,36], we performed a follow-up analysis directly comparing on-beat and off-beat amplitudes at Cz. The amplitude of the P3 significantly differed between on-beat and off-beat trials at Cz (*F*(1,35) = 5.25, *p* = .03, η^2^_p_ = .13). In contrast to the effects of stimulus timing on the N2 and P3, the timing of stimulus presentation did not influence the amplitude of the LPC (*F*(1,35) = 1.80, *p* = .19, η^2^_p_ = .05), although this effect approached significance (*p* = 0.07) when analysis was restricted to electrode Cz where amplitude differences appeared to be the greatest. There was a main effect of electrode for the LPC (*F*(1,35) = 80.05, *p* < .001, η^2^_p_ =.70), but no timing x electrode interaction (*F*(1,35) = 1.34, *p* = .25, η^2^_p_ = .04).

#### Rhythmic modulation of memory (RMM) effects

The above results reveal that background auditory rhythms influence post-perceptual processing as indexed by the N2 and P3 ERPs. To more directly test the mnemonic relevance of these effects, we next compared the amplitude of these components for on-beat and off-beat trials within each participant group (RMM, No-RMM) (Fig 3). For the N2 component, ERP amplitudes were enhanced for on-beat compared off-beat trials in participants who demonstrated a rhythmic modulation of memory effect (RMM group; *F*(1,21) = 9.52, *p* = .006, η^2^_p_ = 0.31). In contrast, there was not a significant difference between on-beat and off-beat amplitudes in participants who did not demonstrate a rhythmic modulation of memory effect (No-RMM group; *F*(1,13) = 1.32, *p* = .27, η^2^_p_ = .09). A similar pattern was present for the P3 component: ERP amplitudes were enhanced for on-beat compared off-beat trials in participants who demonstrated a rhythmic modulation of memory effect (RMM group; *F*(1,21) = 5.78, *p* = .03, η^2^_p_ = .22) while ERP amplitudes for on-beat and off-beat trials did not differ in participants who did not demonstrate a rhythmic modulation of memory effect (No-RMM group; *F*(1,13) = .08, *p* = .79, η^2^_p_ = .006).

We next investigated whether stimulus-evoked responses differed for the RMM and No-RMM groups regardless of stimulus timing (Fig 4). Mean amplitudes for each component were entered into separate mixed ANOVAs with within-subjects factors of electrode and between-subjects factors of group (RMM, no-RMM). For the early N1 component, there was no main effect of group (RMM, no-RMM) nor group x electrode interaction (*Fs* < 1). This was also the case for the N2 and P3 components. While the main effect of group also did not reach significance for the LPC (*F*(1,34) = 2.46, *p* = .12 η^2^_p_ = .10), visual inspection of the topography of the effect suggested that the difference between groups had a more frontal distribution (Fig 4B). Given that ERP subsequent memory effects are often greatest over more frontal electrode sites [44,45], we performed a follow-up analysis restricted to the frontal electrode site (Fz). Amplitudes significantly differed between the two memory groups at Fz (*t*(34) = 2.11, *p* = .04, d = .72), with reduced amplitudes in the RMM group compared to the no-RMM group.

**Figure 4.**
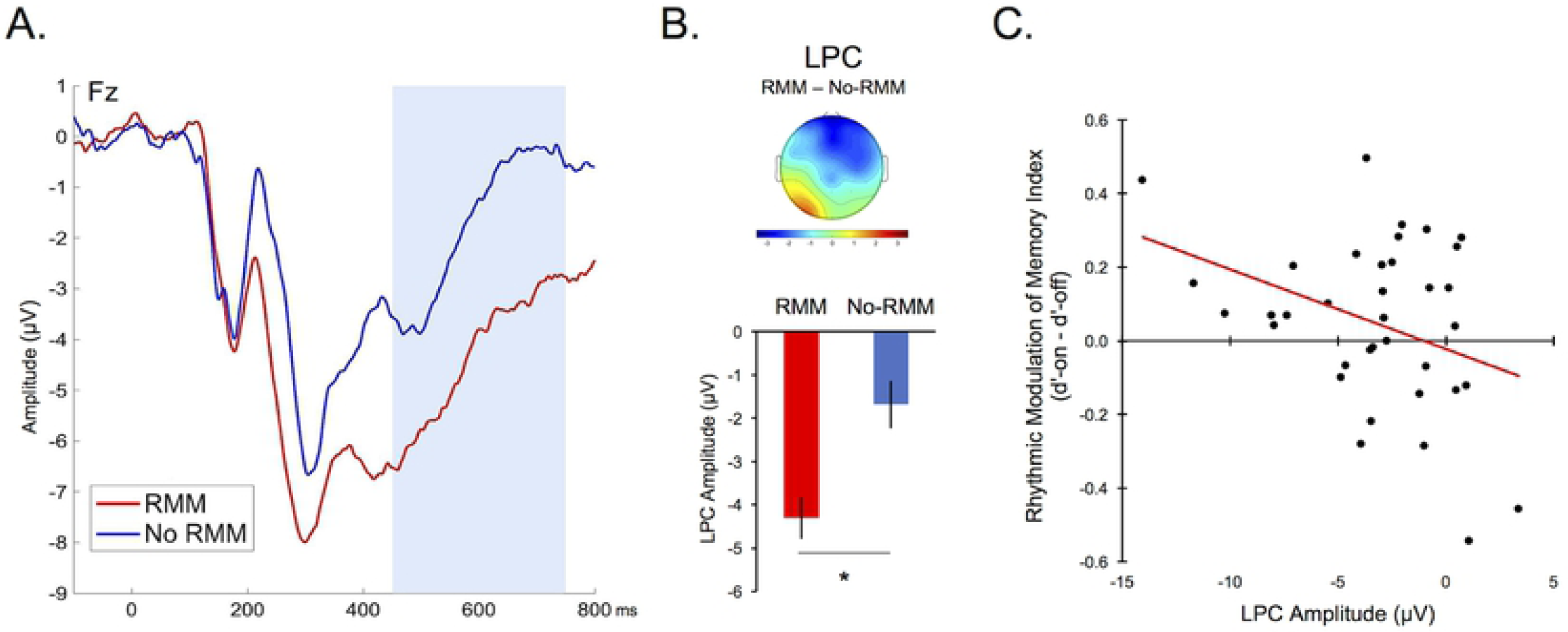
Rhythmic modulation of memory effect. (A) ERP waveform for each memory group (RMM, No-RMM) time-locked to the onset of the visual stimulus at representative electrode Fz. (B) Scalp topography of the average LPC difference between RMM and no-RMM groups during the blue-shaded time window in panel A; bar chart shows average LPC amplitude at electrode Fz for each memory group during the LPC. Bar plots depict average signal for each group. LPC amplitude was significantly reduced in the RMM group compared with the no RMM group. (C) LPC amplitude at electrode Fz was negatively correlated with the rhythmic modulation of memory index (memory on-beat – memory off-beat). Error bars = SEM. * p < .05.

#### Brain-Behavior Correlations

Following the approach used in prior studies [7,22], we next conducted between-subjects correlations to investigate the relationship between stimulus-evoked neural responses and individual differences in the effects of rhythm on memory performance. First, we examined the post-perceptual N2 and P3 components which showed a significant effect of stimulus timing, particularly at electrode Cz. There was no correlation between the rhythmic modulation of memory index (d’-on minus d’-off) and amplitude differences (on-beat minus off-beat trials) across individuals for the N2 or the P3 (*p*s > .20). Next, we examined the later frontal positivity which showed a significant effect of group (RMM, No-RMM) at electrode Fz, where the effect of group was greatest. There was a significant negative correlation between the rhythmic modulation of memory index and the LPC amplitude at Fz (*r*(36) = −.36, *p* = .03) (Fig 4C).

We then conducted two sequential multiple regression analyses to explore whether individual differences in earlier neural responses to rhythm (N2/P3 modulations for on-beat vs. off-beat trials) interact with the later ERP effects (LPC) to predict behavioral performance. Predictors and outcome variables were all linearly related and followed a normal distribution. No multivariate outliers were present in either model and there were no extreme univariate outliers (>3SD from the mean) present in the outcome variable or any of the predictors. In the first model, the N2 effect (amplitude difference for on-beat vs. off-beat trials at electrode site Cz) and the LPC effect (mean amplitude at electrode site Fz) were entered as continuous predictors with the rhythmic modulation of memory index (subsequent memory for on-beat minus off-beat trials) as the dependent variable. Together, the N2 and LPC predictors did not account for a significant amount of variance in memory performance (*R*^*2*^ = .13; *F*(3,32) = 2.46, *p* = .10), although LPC amplitude still individually predicted RMM (*B* = −.02, *β* = −.35, *p* = .04). However, the addition of the interaction term in the second step significantly improved the model *(R*^*2*^ *change* = .11, *F*(1,32) = 4.45, *p* = .04) such that it accounted for a significant amount of variation in behavioral performance (*R*^*2*^ = .24; *F*(3,32) = 3.29, *p* = .03). Importantly, while neither the N2 (*B* = .03, *β* = .24, *p* = .26) nor the LPC (*B* < .001, *β* = .002, *p* = .99) were reliable predictors of the rhythmic modulation of memory in this model, the interaction term was significant (*B* = .02, *β* = .58, *p* = .04). This reveals that individual differences in participants’ neural response to rhythm earlier in the processing stream moderates the relationship between the LPC amplitude and the rhythmic modulation of memory. Specifically, individuals who had the greatest N2 difference for on-beat and off-beat trials (e.g., greatest attentional orienting effect) and the lowest LPC amplitudes (e.g., potentially an index of reduced higher-level cognitive processing) demonstrated the largest effect of rhythm on subsequent memory performance (Fig 5). The second multiple regression analysis mirrored the first analysis, with the only difference being that the P3 effect (amplitude difference for on-beat vs. off-beat trials at Cz) was entered as a continuous predictor rather than the N2 effect. This model did not account for a significant amount of variance in memory performance regardless of whether or not the interaction term was included (*p*s > .09), suggesting that the P3 effect did not moderate the relationship between the LPC and the rhythmic modulation of memory. Post analysis diagnostics of both regression models confirmed the reliability of our results, as there were no influential cases present (Cook’s Distance < 1), the assumption of homoscedasticity was met, and multicollinearity between predictors was not present (Tolerance > .10; VIF < 10).

**Fig 5.**
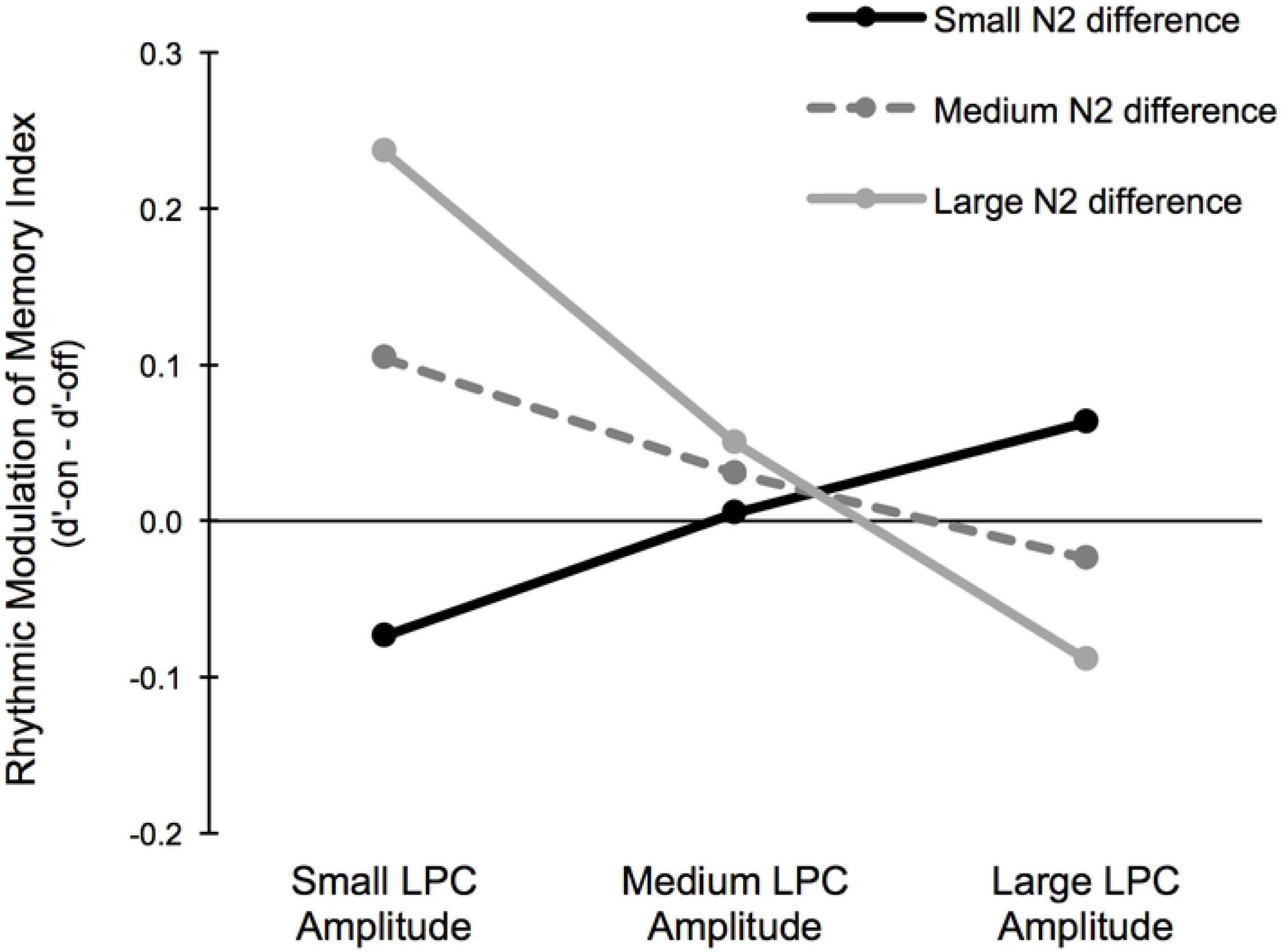
Relationship between post-perceptual ERPs and effects of rhythm on subsequent memory. The relationship between LPC amplitudes and the rhythmic modulation of memory (better memory for on-beat versus off-beat trials) is moderated by individual differences in participants’ neural response to rhythm earlier in the processing stream (N2). Given that LPC amplitudes were significantly different for the RMM and no RMM group at electrode Fz, the LPC amplitudes in the regression analysis were measured at electrode Fz. Since differences in the N2 were seen at electrode Cz, the N2 effect (amplitude difference for on-beat – off-beat trials) is measured at electrode Cz. Small, medium, and large bins for the LPC and N2 effects were defined based on the SD of each predictor. Rhythmic modulation of memory index = d’-On – d’-Off.

## Discussion

The current study provides novel electrophysiological evidence that environmental rhythms influence long-term memory formation by guiding attention to specific moments in time. During incidental encoding, stimulus-evoked neural responses differed for visual images that appeared in synchrony versus out-of-synchrony with a background auditory beat. Amplitude differences were present in post-perceptual ERP components (e.g., N2, P3) associated with temporal attention and were present in participants demonstrating rhythmic modulation of memory (superior subsequent memory for on-beat compared to off-beat trials). The behavioral effect of rhythm on memory performance was also related to the amplitude of a later frontal positivity during the time window of classic subsequent memory effects. Together, these results support the proposal that rhythm directs domain-general attentional resources to specific moments in time [11], and suggests that the dynamic effects of rhythm on memory encoding reflect post-perceptual attentional selection as well as downstream changes in higher-order cognitive processing.

A key finding of the current study was that the timing of visual stimulus presentation during encoding (on-beat vs. off-beat) influenced the amplitude of later ERP components (e.g., N2, P3) rather than earlier ERP components (e.g., N1). This finding is consistent with prior work demonstrating effects of temporal attention on visually-evoked N2 and P3 components [30,36,37], and reveals that temporal cues provided by musical rhythm influence post-perceptual visual processing. While rhythmic temporal cues have also been shown to influence the amplitude of earlier N1 components associated with initial stimulus processing [7,27], the modulation of sensory/perceptual ERPs is not always observed following manipulations of temporal attention and may depend on the nature of the temporal cues or task context [19,38-42]. For example, in prior studies demonstrating rhythmic effects on the N1, participants performed perceptually-demanding target detection/discrimination tasks [7,27]. Correa and colleagues have argued that while later ERPs (e.g., N2, P3) are consistently modulated by temporal attention, early ERPs (e.g., N1) are only affected by temporal orienting cues in the context of tasks that place high demands on perceptual processing (e.g., target discrimination) [30]. In the current study, perceptual demands were relatively low, as participants were tasked with making a semantic decision about each stimulus as it appeared. Thus, it is possible that earlier effects could emerge in a more perceptually-demanding task. Importantly, our finding that rhythmic temporal cues influence stimulus processing at relatively late stages of processing suggests that in addition to acting as a mechanism of early attentional selection [1,2,14,15,50,51], rhythmic temporal cues can also act as a mechanism of selection at post-perceptual stages of processing.

In prior studies, effects of temporal attention on the N2 and P3 components have been interpreted as reflecting attention-related modulations of decision or response-related processing [30,34,36,37]. While the direction of the amplitude modulation of the N2 and P3 for temporally-predicted versus temporally-unpredicted stimuli differs across studies, amplitude modulations like those observed in the current study (increased N2 amplitudes and decreased P3 amplitudes) have been observed when stimuli appear at attended locations/times or are contextually expected (i.e., not deviant) [35,52-56]. In the current study, amplitude modulations of the N2 and P3 could therefore reflect the tuning or sharpening of decision-related or response-related processes which optimize the encoding of rhythmically-predicted stimuli that appear in synchrony with the beat (e.g., serve as a temporal filter) [57,58]. Support for this proposal comes from the finding that in addition to being better remembered, on-beat stimuli in the current study were also more rapidly classified during incidental encoding compared to off-beat stimuli. Furthermore, post-hoc analysis indicates that while the amplitude of the N2/P3 did not correlate with overall response times during encoding, there was a significant correlation between P3 amplitude at electrode Cz and the response facilitation for on-beat versus off-beat stimuli at encoding (*r*(36) = −.34, *p* = .04), whereby participants demonstrating faster classification times for on-beat versus off-beat stimuli during encoding also demonstrated greater P3 amplitude reductions. Though the functional significance of the N2/P3 amplitude modulations represent an important area for future research, these results are congruent with the proposal that rhythmic modulation of mid-latency components reflects facilitated decision or response-related processing.

A more direct link between rhythmic effects on post-perceptual stimulus processing and memory encoding comes from the finding that the N2 and P3 amplitudes not only differed according to the timing of stimulus presentation within the rhythmic stream, but also differed according to whether or not participants demonstrated an effect of stimulus timing on subsequent memory. Specifically, a significant difference in N2 and P3 amplitudes for on-beat versus off-beat stimuli was only present in participants who demonstrated a positive rhythmic modulation of memory effect (better subsequent memory for on-beat versus off-beat stimuli). This suggests that the way in which temporally-predicted (on-beat) versus temporally-unpredicted (off-beat) stimuli are initially processed impacts how well they are later remembered. However, this effect of rhythm on post-perceptual ERPs was not directly associated with the effect of rhythm on later memory performance, as neither the N2 nor P3 modulations correlated with the mnemonic benefit for on-beat versus off-beat stimuli. Instead, the rhythmic effects on attention-related components (specifically the N2) interacted with amplitude modulations of the more positive late positive complex (LPC) over frontal electrodes to predict memory performance. This raises the intriguing possibility that the effect of rhythm on memory encoding reflects an interaction between earlier attentional selection and downstream cognitive processing.

In contrast to the N2 and P3 ERP effects, the timing of stimulus presentation (on-beat vs. off-beat) within the rhythmic stream did not influence the amplitude of the LPC. Instead, the amplitude of the LPC differed according to whether or not participants demonstrated better subsequent memory for on-beat vs. off-beat stimuli, with reduced LPC amplitudes in participants demonstrating greater rhythmic modulation of memory. The directionality of this amplitude difference may initially seem at odds with the typical finding in the subsequent memory literature of greater LPC amplitudes for items that are later remembered versus forgotten. However, the direction of the ERP subsequent memory effect can differ according to the nature of the to-be-remembered stimulus (e.g., words vs. non-words) [45]. Additionally, it is important to note that in the current study LPC modulations do not reflect subsequent memory performance per se (i.e., whether individual stimuli will be later remembered or forgotten) but rather reflect individual differences in the *effect of rhythm* on subsequent memory performance. Thus, rather than serving as an index of general encoding success, modulations of the LPC may reflect the degree to which higher-level cognitive operations are required for stimulus processing later in the processing stream. Prior studies have found that the LPC is modulated by higher-level control demands [59-61]. In the context of the current study, demands on controlled processing may be reduced when environmental rhythms can be leveraged to direct attention in time. For example, greater LPC amplitudes in the current study could reflect more intensive or prolonged semantic processing in participants who are less able to leverage rhythmic temporal cues to enhance memory encoding. This interpretation aligns with the finding that individual differences in earlier ERP effects of temporal orienting (e.g., N2 amplitude difference for on-beat versus off-beat trials) moderated the relationship between the amplitude of the LPC and the rhythmic modulation of memory. Additional support for this interpretation comes from the observation that the amplitude of the LPC negatively correlated with the rhythmic modulation of memory index across participants. Interestingly, Jones and Ward recently observed a similar result, whereby LPC modulations were negatively related to the effects of rhythm on memory performance across participants [7]. Although future studies should investigate the specific processes indexed by these late ERP effects, together these results suggest that later stages of processing play an important role in the effects of rhythm on memory performance

While the results of the current study support the hypothesis that external rhythms dynamically modulate memory encoding by influencing post-perceptual stimulus processing, an important outstanding question is whether rhythm also impacts memory by modulating neural activity *before* the onset of a stimulus. Prior research has demonstrated that pre-stimulus neural activity can predict whether or not visual events are effectively encoded into memory and have been interpreted as reflecting the degree to which encoding-related attentional processes are prepared ahead of stimulus presentation [62,63]. Temporal orienting of attention has also been shown to influence anticipatory neural responses before the onset of a stimulus [37,64]. Thus, external rhythms that direct attention to particular moments in time may also influence memory encoding by regulating cortical excitability prior to the onset of a stimulus. To address this possibility, future work should assess how pre-stimulus activity, such as the pre-stimulus phase of entrained oscillations or pre-stimulus alpha/beta power, influences the rhythmic modulation of memory.

A second outstanding question is whether there is a relationship between the evoked neural responses observed in the current study and previously reported effects of rhythm on oscillatory entrainment. Using the same paradigm, Hickey and colleagues recently demonstrated that rhythmic effects on memory encoding are closely related to how strongly rhythm is represented in the brain (i.e., oscillatory entrainment to the rhythmic beat) [22]. Although it is important to note that the ERPs in the current study were measured in response to the onset of the to-be-remembered visual stimuli rather than the onset of background auditory beat, and are therefore unlikely to be measuring the same type of neural response captured by entrainment measures, these two types of neural responses could be related and together may serve to better predict the effect of rhythm on memory encoding [16,65-68]. Future research should aim to clarify how these different types of neural responses to rhythm have independent and/or interactive effects on memory encoding [54,66].

